# Revealing new therapeutic opportunities through drug target prediction via class imbalance-tolerant machine learning

**DOI:** 10.1101/572420

**Authors:** Siqi Liang, Haiyuan Yu

**Affiliations:** Department of Biological Statistics and Computational Biology, Cornell University, Ithaca, New York, 14853, USA; Weill Institute for Cell and Molecular Biology, Cornell University, New York, 14853, USA

## Abstract

*In silico* drug target prediction provides valuable information for drug repurposing, understanding of side effects as well as expansion of the druggable genome. In particular, discovery of actionable drug targets is critical to developing targeted therapies for diseases. Here, we develop a robust method for drug target prediction by leveraging a class imbalance-tolerant machine learning framework with a novel training scheme. We incorporate novel features, including drug-gene phenotype similarity and gene expression profile similarity, that capture information orthogonal to other features. We show that our classifier achieves robust performance and is able to predict gene targets for new drugs as well as drugs that target unexplored genes. By providing newly predicted drug-target associations, we uncover novel opportunities of drug repurposing that may benefit cancer treatment through action on either known drug targets or currently undrugged genes.

## INTRODUCTION

Target identification is a crucial step during drug development. As the cost of bringing a single new drug to market skyrockets to over 2.7 billion dollars on average [1], alternative approaches, such as drug repurposing, have been pursued with increasing efforts. For example, the drug aspirin, commonly used for treating fever and acute pain, has been found in recent years to show anti-cancer activities through attenuation of EGFR expression [2], inhibition of COX-2 [3] and suppression of NF-κB activation by TNF [4]. As a result, the efficacy of aspirin in treating multiple types of cancers, including breast cancer, prostate cancer and colorectal cancer, is being actively evaluated in clinical trials. By repurposing approved drugs for new indications through novel target discovery, the cost of drug development can be substantially reduced, especially in the preclinical and earlier clinical phases where the toxicity and dosage of the drug is assessed [5]. In addition to benefiting drug repurposing efforts, identifying unknown targets of drugs can facilitate understanding of their side effects, which are often caused by drugs binding to unintended targets. The serotonin receptor agonist cisapride, as an example, is a gastroprokinetic agent used for treating gastric reflux, but it can cause serious cardiac events including arrhythmia and even lead to death. The mechanism behind the cardiac effects of cisapride was discovered in 1997 to be its high-affinity blocking of the human cardiac potassium channel [6]. And this resulted in its withdrawal from the US market three years later. Furthermore, out of over 4,400 genes estimated to be druggable in the human genome [7], only less than half of them are currently targeted by approved drugs. Therefore, identification of novel gene targets can help with expanding the druggable genome, opening up new avenues for drug development.

Experimental methods for determining drug-target associations provides direct evidence and information on the mode of action of drugs. However, their high cost and long timeframe have prohibited them from large-scale application. As an alternative, computational approaches, including docking-based methods and machine learning-based methods, have been developed to predict new drug-target associations [8]. In particular, machine learning-based methods that exploit the chemogenomic space have yielded considerable success in drug target prediction without requiring three-dimensional protein structures of the targets [9-13]. Various features, including chemical similarity [14, 15] and side effect similarity [16, 17], have proved valuable in identifying new associations between drugs and targets. Nevertheless, two fallacies are commonly overlooked: conventional train-test splitting and cross-validation schemes are flawed for pair-input prediction tasks [18]; extreme class imbalance in drug target datasets is not satisfactorily addressed by commonly used methods such as sampling from the majority class [19]. Moreover, most methods lack the ability to predict drug-target interactions for genes that are not yet known to be druggable.

To address these challenges, in this study, we design a novel training scheme that prevents possible overfitting caused by overlapping drugs or targets in the training and test sets and at the same time solves the class imbalance problem with an ensemble method. Additionally, we exploit two new types of features, namely the phenotype similarity between a drug and a gene, and the expression profile similarity between two genes across different tissues. We show that they confer considerable predictive power and provide orthogonal information that is not captured by other features. Incorporating these features, we build a classifier and demonstrate that it achieves robust performance. Further, our classifier is able to make predictions for drugs without known targets and for genes that are not yet known to be druggable. By predicting new potential drug-target associations, we reveal unexplored opportunities of drug discovery and repurposing for cancer treatment.

## RESULTS

### Drug-gene phenotype similarity and gene expression profile similarity provides complementary information for identifying drug targets

Similarity-based features have been widely used for drug target prediction [20]. Behind them is a simple motivating hypothesis: similar drugs tend to have the same gene targets, and correspondingly, similar genes tend to be targeted by the same drugs. Among various drug-drug similarity metrics, chemical similarity and side effect similarity have been most extensively employed [14-17]. We obtained a comprehensive dataset of known drug-target associations by extracting relevant information for all drugs with human gene targets from a recent version of the Probes & Drugs database [21]. Further filtering (see Methods) resulted in a total of 1,262 drugs with 11,556 drug-target associations involving 1,062 human genes.

To calculate drug-drug similarity features, for each drug-gene instance, we considered all drugs that are known to target the gene in question and measured their resemblance to the drug in question in terms of chemical similarity and side effect similarity. Since a gene could have multiple known targeters, different aggregation functions were applied to obtain real-valued features for each drug-gene pair (Fig. 1a). 2D chemical similarity between two compounds was calculated by taking the Jaccard index of their Morgan fingerprints, which represent planar chemical substructures in the form of bit vectors [22]. Not surprisingly, when taking maximum, mean or median as the aggregation function, drugs are significantly more chemically similar to known targeters of their gene target than to known targeters of other genes (Fig. 1b). On a similar note, computing similarity by taking the Jaccard index of their side effects (see Methods) gave identical trends (Fig. 1c). But interestingly, when aggregating similarity scores by the minimum, drug-gene pairs that are known to be associated had significantly lower scores than those that are not known to be associated, regardless of the type of similarity metric used (Fig. 1b, 1c). This can be explained by the fact that genes in associated drug-gene pairs have a significantly higher number of known targeters in a broader chemogenomic space than genes in other drug-gene pairs (Supplementary Fig. 1a). Recently, a method for encoding the 3D structure of molecules has been developed and has been shown to enhance the performance of conventional 2D fingerprinting methods in binding prediction [23]. Using the Jaccard index of the 3D molecular fingerprints as the chemical similarity metric, we discovered similar trends as using 2D chemical similarity and side effect similarity (Fig. 1d). Notably, 3D chemical similarity features are only weakly correlated with 2D chemical similarity and side effect similarity features (Supplementary Fig. 1b), providing new information about the relatedness of two drugs.

**Figure 1.**
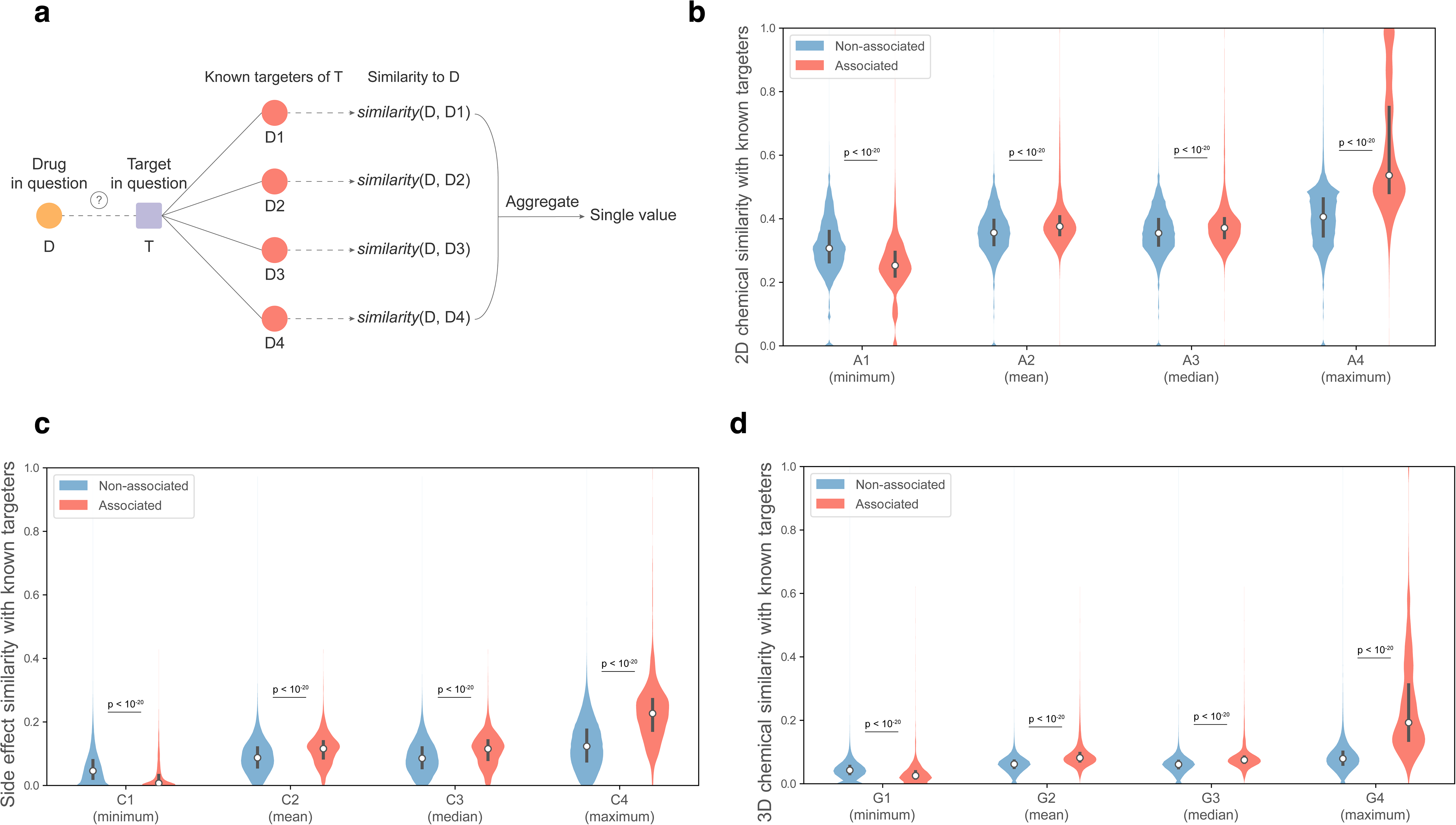
Drug-drug similarity features. (a) Schematics of calculating drug-drug similarity features for each drug-gene pair. Each group of features consists of four features corresponding to four different aggregation functions: minimum, mean, median and maximum. (b) Distribution of 2D chemical similarity features (feature group A). (c) Distribution of side effect similarity features (feature group C). (d) Distribution of 3D chemical similarity features (feature group G). (Statistical significance determined by the two-sided Mann-Whitney U test)

In addition to aforementioned feature groups, which have already been incorporated in previous drug-target prediction methods, here we introduce two novel types of features: drug-gene phenotype similarity and expression profile similarity between two genes. Drugs that act directly on a protein and alter its activity may lead to the same phenotypic changes as mutations on the corresponding gene. On this account, we designed a drug-gene phenotype similarity metric by taking the Jaccard index of the side effects of the drugs and disease phenotypes of the gene (Fig. 2a). As expected, drug-gene pairs that are known to be associated have significantly higher phenotype similarity scores than drug-gene pairs that are not known to be associated (Fig. 2b). On top of drug-drug and drug-gene similarity features, we calculated similarity between two genes as their correlation coefficient in expression levels across different tissues using gene expression data from GTEx [24]. To obtain scalar features, we considered the similarity between the gene in question and known targets of the drug in question and applied the same four aggregation functions as drug-drug similarity features (Fig. 2c). We discovered that when taking maximum, mean or median as the aggregation function, genes have significantly more similar expression profiles to known targets of their targeters than to known targets of other drugs (Fig. 2d). Using minimum as the aggregation function rendered the opposite trend, which could be explained by drugs in drug-gene pairs that are known to be associated having a significantly more diverse target set than drugs in other drug-gene pairs (Supplementary Fig. 1c). Intriguingly, expression profile features, especially when aggregated with maximum, mean or median, exhibit almost no correlation with other groups of features, bringing in complementary information that other features do not capture (Fig. 2e).

**Figure 2.**
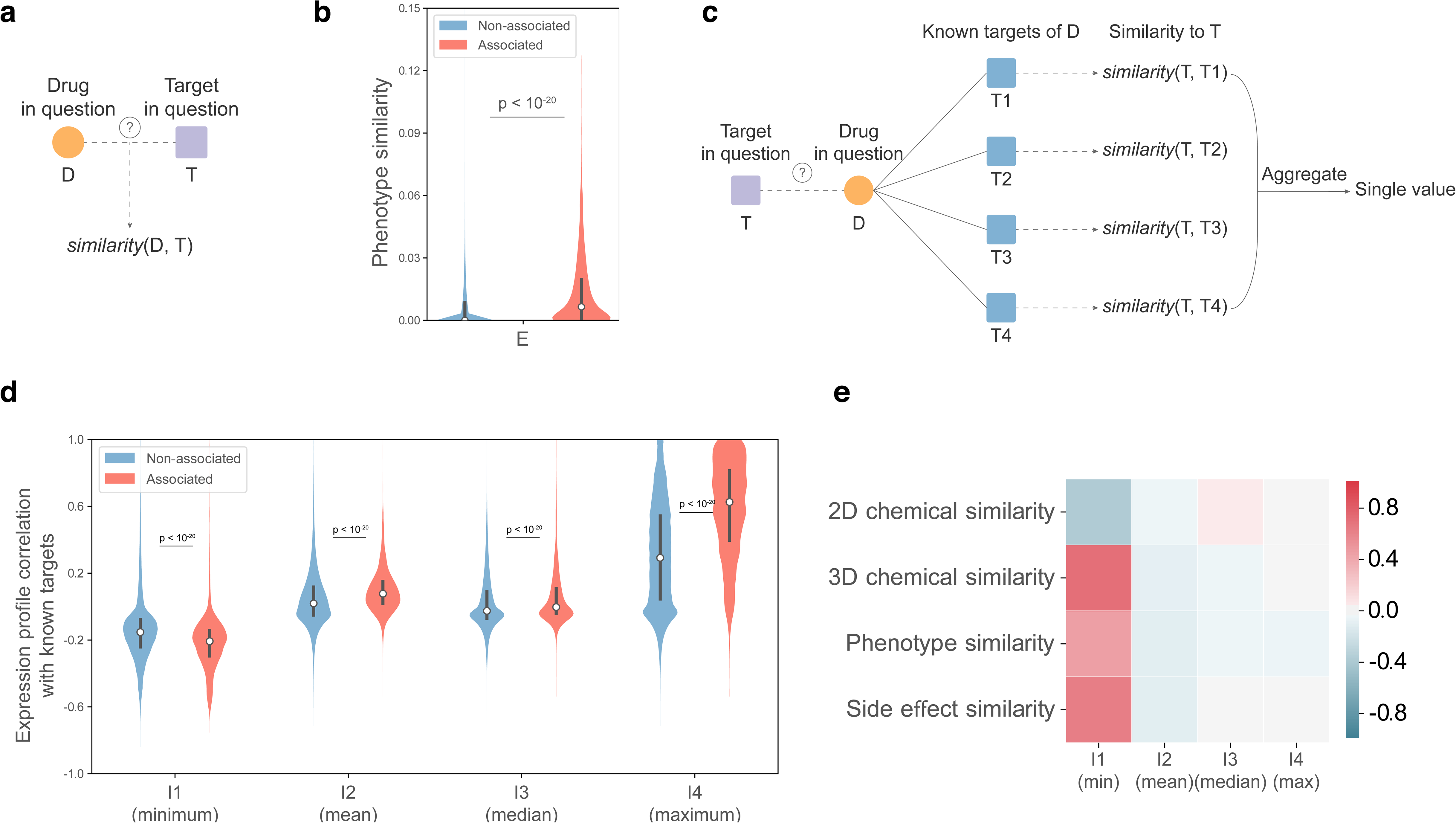
Drug-gene phenotype similarity and gene expression profile similarity features. (a) Schematics of calculating the drug-gene phenotype similarity feature. (b) Distribution of the drug-gene phenotype similarity feature (feature group E). (c) Schematics of calculating gene expression profile similarity features. This group of features consists of four features corresponding to four different aggregation functions: minimum, mean, median and maximum. (d) Distribution of gene expression profile similarity features (feature group I). (e) Spearman correlation coefficients of gene expression similarity features with other types of features. Only the correlation coefficient corresponding to the most correlated or anti-correlated feature in each type of features is shown. (Statistical significance determined by the two-sided Mann-Whitney U test)

It is worth noticing that drug-gene phenotype similarity and gene expression profile similarity features can be calculated even if the gene in question has no known drugs that targets it. This potentializes us to make predictions for currently “undrugged” genes, thereby expanding the druggable genome. To extend this advantage to drug-drug similarity features, we considered targeters of protein-protein interaction partners of the gene in question for both chemical similarity and side effect similarity (Supplementary Fig. 2a). We also considered protein-protein interactors of the gene in question for drug-gene phenotype similarity (Supplementary Fig. 2b). Four new groups of features were thus generated, and we showed that they possess distinguishing power in separating drug-target pairs and other drug-gene pairs (Supplementary Fig. 2c-2f).

### A novel training scheme prevents overfitting and solves the class imbalance problem

In order to build a machine learning model for drug-target prediction, we divided all drug-gene pairs into a training set and a test set. If the split is random, the machine learning algorithm might pick up characteristics of single drugs or genes that appear in both the training set and the test set, causing a problem called overfitting. To avoid this, we applied a splitting scheme where the drugs were first randomly divided into “train drugs” and “test drugs”, and the genes were split into “train targets” and “test targets” (Fig. 3a), so that there is no overlap between the training set and the test set in terms of either drugs or genes. Since there was no gold-standard dataset of non-associated drug-gene pairs, all drug-gene pairs not known to be associated were considered as non-associated. This resulted in an extreme class imbalance where negative instances were over 100 folds more than positive instances in quantity. To address this problem, we divided the negative instances (non-associated pairs) in the training set into a number of subsets, and each subset was combined with all the positive instances (associated pairs) in the training set to obtain a training subset (Fig. 3b). For every training subset we trained a classifier, and eventually we would take the average prediction score of the ensemble of classifiers as our final prediction score.

**Figure 3.**
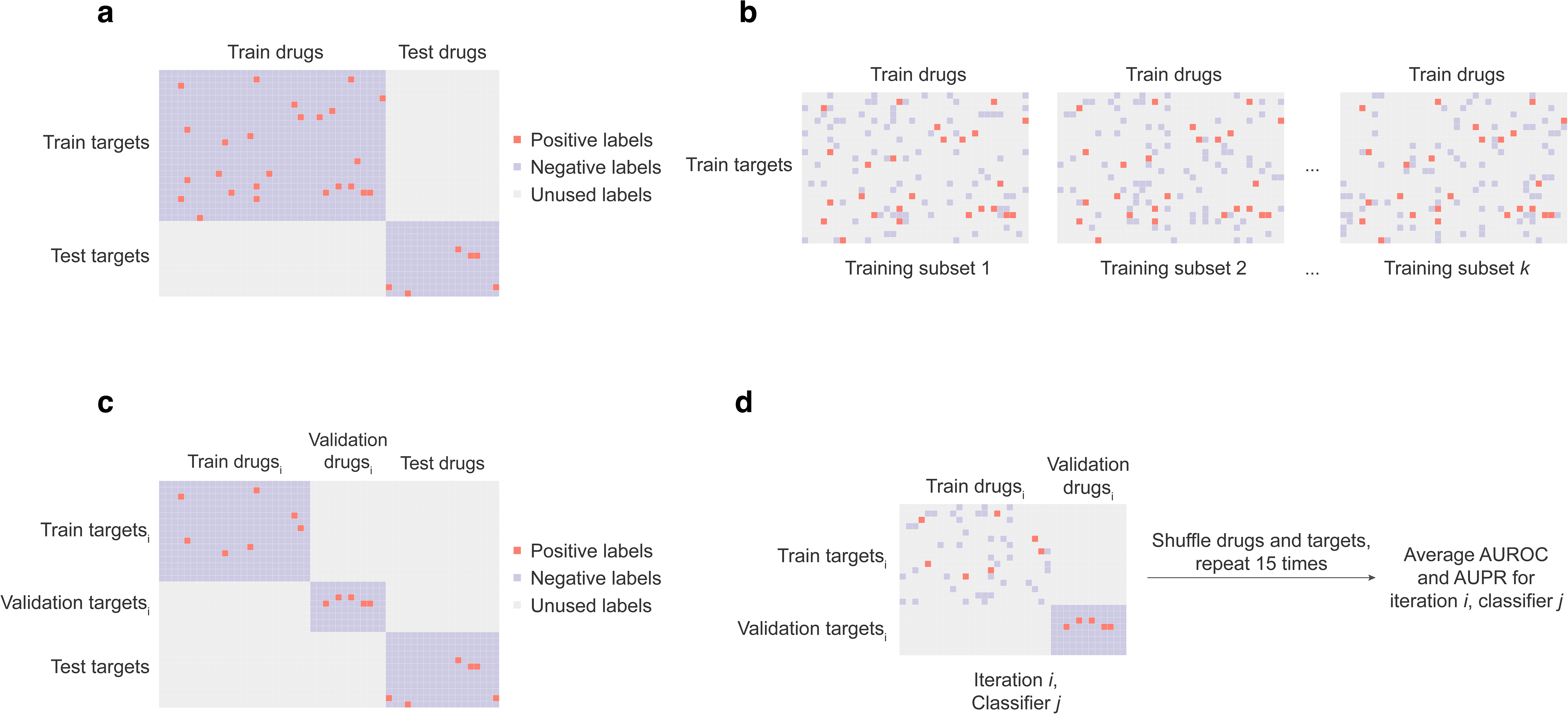
Data splitting and the training scheme. (a) The train-test split. All drugs were split into “train drugs” and “test drugs”, while all genes were split into “train targets” and “test targets”. The training set then consisted of all drug-gene pairs where the drug was a “train drug” and the target was a “train target”. Similarly, the test set consisted of all drug-gene pairs where the drug was a “test drug” and the target was a “test target”. (b) The training set was split into multiple training subsets by splitting all the training examples with a negative label into subsets, each combined with all examples with a positive label to form a training subset. (c) The entire training set was split into one portion used for fitting the classifiers and one portion used for validation using a similar splitting method as the train-test split so that there was no overlap between data used for fitting the classifiers and data used for validation in terms of either drugs or genes. (d) For each classifier and each set of hyperparameters, the train-validation split was done 15 times. Each time the training data was intersected with the corresponding training subset before used for classifier fitting, while the entire validation set was used for performance evaluation using AUROC and AUPR as metrics. The average AUROC and AUPR over the 15 splits were considered as the performance for that specific set of hyperparameters.

The use of typical cross-validation could prevent classifiers from achieving robust predictive performance for pair-input data [18]. Here, we designed a novel training scheme where the hold-out validation set has no overlap with data used for fitting the classifier in terms of either drugs or genes by adopting the same splitting method as the train-test split (Fig. 3c). For each classifier and each set of hyperparameters, this splitting was done 15 times, and each time the drug-gene pairs used for model fitting was intersected with the corresponding training subset while all the drug-gene pair used for validation was used for evaluating model performance (Fig. 3d). This training scheme solved the class imbalance problem while utilizing all training instances.

To pick out the most informative set of features and speed up the training process, we adopted a feature selection method known as group minimax concave penalty (MCP), which has been used for biological feature selection previously [25]. The final set of features comprised of 14 features, with every type of features present, including the newly proposed phenotype similarity and expression profile similarity features. Since drug-target and protein-protein networks were exploited for feature calculation, no information about drugs and genes used for evaluating performance was made visible when calculating features for training (Supplementary Fig. 3a). On the other hand, when calculating the feature matrix for performance evaluation, only associations between drugs and genes to be predicted were masked (Supplementary Fig. 3b). The extreme gradient boosting (XGBoost) model [26] was chosen for the classifiers because of its speed and strong performance as shown in recent studies [27, 28]. Using a Bayesian tree-structured Parzen estimator (TPE) approach [29], which has recently been shown to improve classifier performance drastically [30], we optimized hyperparameters for all classifiers in the ensemble (Supplementary Fig. 4a-b). This resulted in an average AUROC of 0.924 across all classifiers in the ensemble and an average AUPR of 0.273 (Fig. 4a). When evaluated on the hold-out test set which has no overlap with the training data in terms of either drugs or genes, we obtained an AUROC of 0.928 (Fig. 4b) and an AUPR of 0.269 (Fig. 4c). This demonstrates that our model was not subject to overfitting and illustrates the robustness of our novel training scheme. Notably, our model attained a precision of 78% on the top 50 predictions and a precision of 48.2% when examining the top 500 predictions. Considering the fact that these are lower bound estimates since drug-gene pairs labeled as non-associated could be actually undiscovered drug-target pairs, our model achieves accurate prediction.

**Figure 4.**
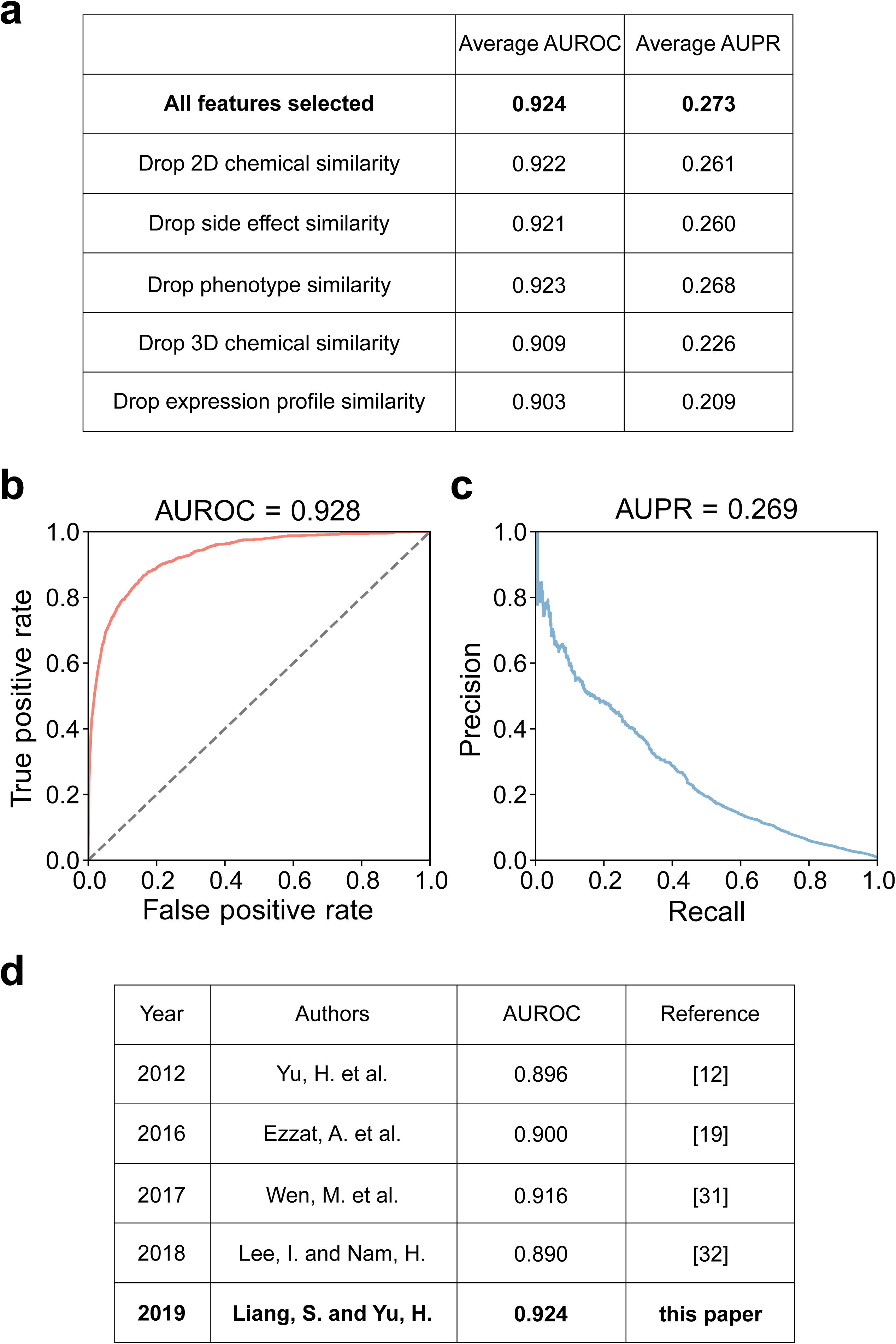
Performance evaluation. (a) Training performance of our model was calculated as the average AUROC and average AUPR across all classifiers in the ensemble using the best sets of hyperparameters. The contribution of each type of features to model performance was evaluated by dropping that type of features from the full model and redoing hyperparameter optimization. Dropping each type of features resulted in a decrease in performance. (b) Receiving operating characteristic (ROC) curve when evaluating the model on the hold-out test set. (c) Precision-recall curve when evaluating the model on the hold-out test set. (d) Comparison of training performance with four recently published methods.

To assess the contribution of each type of features to model performance, we dropped each type of features and retrained our classifier ensemble using the same number of TPE iterations. As expected, model performance declined when any of the 5 groups of features was taken out. Interestingly, exclusion of expression profile similarity resulted in the largest performance drop, followed by exclusion of 3D chemical similarity (Fig. 4a). These results demonstrate the effectiveness of using our newly proposed features for prediction of drug targets. Furthermore, our classifier outperforms other drug-target prediction methods published in recent years [12, 19, 31, 32] (Fig. 4d), achieving favorable performance with a highly imbalanced dataset.

### Newly predicted drug-target associations reveal unexplored opportunities for cancer treatment

We applied our trained model on drug-gene pairs that were not previously known to be associated and predicted novel drug-target associations. By examining newly predicted associations between known drugs and genes that are already known to be druggable (Table 1), we discovered new drug repurposing opportunities. As an example, the antipsychotic drug, fluphenazine, is predicted to target the *PRKDC* gene with high probability. Commonly prescribed for treatment of schizophrenia, fluphenazine primarily acts on dopamine receptors and G-protein coupled receptors [33, 34]. The *PRKDC* gene encodes a DNA-dependent protein kinase (DNA-PKc) that mediates non-homologous end joining (NHEJ) [35], which is an important mechanism by which cells can repair double-strand breaks in DNA without a homologous template [36]. Cells deficient of the *ATM* gene, which is commonly mutated in various types of cancers [37], can evade p53-mediated apoptosis but become reliant on NHEJ for DSB repair [38]. Therefore, while *ATM*-deficient cancers are largely resistant to genotoxic chemotherapy, it has been reported that exposure to a DNA-PKc inhibitor diminishes their ability to repair DSBs and prolongs survival using a *ATM*-deficient mouse lymphoma model [39]. This, along with studies in other cancer types [40, 41], establishes the *PRKDC* gene as a potential target for treatment of *ATM*-deficient cancer and opens up the possibility of repurposing fluphenazine for cancer treatment (Fig. 5a).

**Table 1.** Top 20 drug-target predictions between drugs with known targets and genes that are known to be targeted by drugs.

| CID | cmpdname | iupacname | Target |
| --- | --- | --- | --- |
| 5566 | Trifluoperazine | 10-[3-(4-methylpiperazin-1-yl)propyl]-2-(trifluoromethyl)phenothiazine | DRD1 |
| 11167602 | Regorafenib | 4-[4-[[4-chloro-3-(trifluoromethyl)phenyl]carbamoylamino]-3-fluorophenoxy]-N-methylpyridine-2-carboxamide | TGFBR2 |
| 3372 | Fluphenazine | 2-[4-[3-[2-(trifluoromethyl)phenothiazin-10-yl]propyl]piperazin-1-yl]ethanol | CALY |
| 8226 | Methylergonovine | (6aR,9R)-N-[(2S)-1-hydroxybutan-2-yl]-7-methyl-6,6a,8,9-tetrahydro-4H-indolo[4,3-fg]quinoline-9-carboxamide | ADRA1A |
| 11167602 | Regorafenib | 4-[4-[[4-chloro-3-(trifluoromethyl)phenyl]carbamoylamino]-3-fluorophenoxy]-N-methylpyridine-2-carboxamide | EPHA1 |
| 443884 | Ergonovine | (6aR,9R)-N-[(2S)-1-hydroxypropan-2-yl]-7-methyl-6,6a,8,9-tetrahydro-4H-indolo[4,3-fg]quinoline-9-carboxamide | CYP2D6 |
| 115237 | Paliperidone | 3-[2-[4-(6-fluoro-1,2-benzoxazol-3-yl)piperidin-1-yl]ethyl]-9-hydroxy-2-methyl-6,7,8,9-tetrahydropyrido[1,2-a]pyrimidin-4-one | HTR5A |
| 8226 | Methylergonovine | (6aR,9R)-N-[(2S)-1-hydroxybutan-2-yl]-7-methyl-6,6a,8,9-tetrahydro-4H-indolo[4,3-fg]quinoline-9-carboxamide | CYP3A4 |
| 5073 | Risperidone | 3-[2-[4-(6-fluoro-1,2-benzoxazol-3-yl)piperidin-1-yl]ethyl]-2-methyl-6,7,8,9-tetrahydropyrido[1,2-a]pyrimidin-4-one | KCND3 |
| 3372 | Fluphenazine | 2-[4-[3-[2-(trifluoromethyl)phenothiazin-10-yl]propyl]piperazin-1-yl]ethanol | PRKDC |
| 3001028 | Estrone sulfate | [(8R,9S,13S,14S)-13-methyl-17-oxo-7,8,9,11,12,14,15,16-octahydro-6H-cyclopenta[a]phenanthren-3-yl] hydrogen sulfate | SERPINA6 |
| 2118 | Alprazolam | 8-chloro-1-methyl-6-phenyl-4H-[1,2,4]triazolo[4,3-a][1,4]benzodiazepine | GABRB2 |
| 115237 | Paliperidone | 3-[2-[4-(6-fluoro-1,2-benzoxazol-3-yl)piperidin-1-yl]ethyl]-9-hydroxy-2-methyl-6,7,8,9-tetrahydropyrido[1,2-a]pyrimidin-4-one | HTR6 |
| 4914 | Procaine | 2-(diethylamino)ethyl 4-aminobenzoate | SCN5A |
| 2519 | Caffeine | 1,3,7-trimethylpurine-2,6-dione | LMNA |
| 4917 | Prochlorperazine | 2-chloro-10-[3-(4-methylpiperazin-1-yl)propyl]phenothiazine | CALM1 |
| 31703 | Doxorubicin | (7S,9S)-7-[(2R,4S,5S,6S)-4-amino-5-hydroxy-6-methyloxan-2-yl]oxy-6,9,11-trihydroxy-9-(2-hydroxyacetyl)-4-methoxy-8,10-dihydro-7H-tetracene-5,12-dione | TP53 |
| 10531 | Dihydroergotamine | (5'α)-9,10-dihydro-12'-hydroxy-2'-methyl-5'-(phenylmethyl)-ergotaman-3',6',18-trione | HTR1A |
| 3696 | Imipramine | 3-(5,6-dihydrobenzo[b][1]benzazepin-11-yl)-N,N-dimethylpropan-1-amine | CYP3A4 |
| 31703 | Doxorubicin | (7S,9S)-7-[(2R,4S,5S,6S)-4-amino-5-hydroxy-6-methyloxan-2-yl]oxy-6,9,11-trihydroxy-9-(2-hydroxyacetyl)-4-methoxy-8,10-dihydro-7H-tetracene-5,12-dione | CYP1A2 |

**Figure 5.**
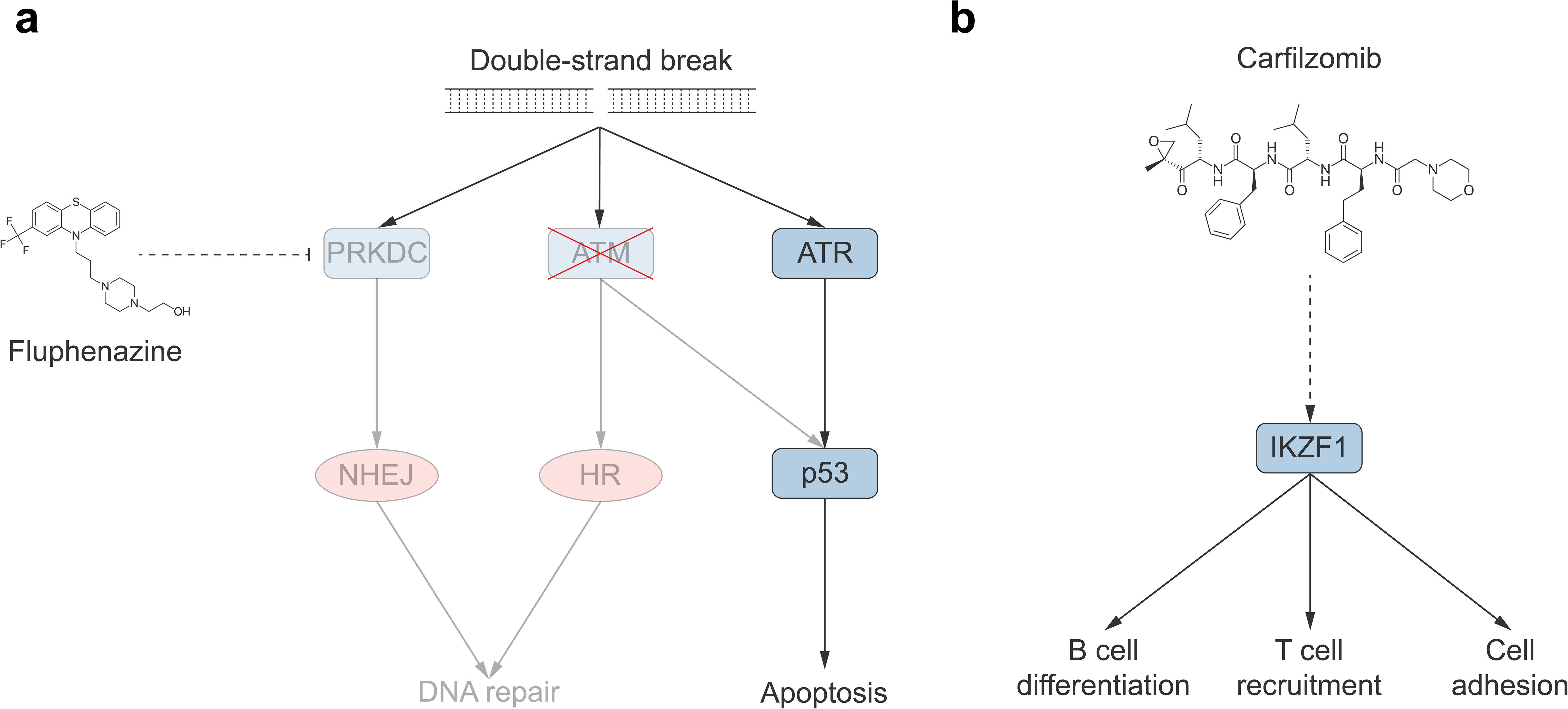
Novel therapeutic opportunities for cancer uncovered by drug target predictions. (a) Fluphenazine is predicted to target DNA-PKc encoded by the *PRKDC* gene. DNA-PKc is a key mediator of non-homologous end joining (NHEJ), which is an alternative mechanism for DNA double-strand break (DSB) repair to homologous recombination (HR). Fluphenazine can potentially be repurposed to treat *ATM*-deficient cancer by disabling NHEJ. (b) Carfilzomib is predicted to target the transcription factor *IKZF1*, which is a modulator of immune responses. Activation of *IKZF1* enhances cell adhesion and promotes B cell differentiation and T cell recruitment. This renders IKZF1 a potential drug target that might enhance the efficacy of immunotherapy when activated.

In addition to predicting associations between known drugs and genes that are already known as drug targets, we also predicted potential associations between drugs and genes that are not yet known to be druggable, in an effort to expand the druggable genome (Table 2). For instance, carfilzomib and *IKZF1* is the drug-target pair with the highest probability to be associated where the gene is not yet known to be druggable. *IKZF1* encodes a transcription factor and has been shown to have tumor suppressive function during leukemia development [42]. It is a modulator of immune responses and its activation facilitates recruitment of T cells (Fig. 5b). Furthermore, according to a recent publication, overexpression of *IKZF1* in tumors results in significantly improved responses to immunotherapy, including anti-PD1 and anti-CTLA4 treatment [43]. These discoveries have rendered *IKZF1* an ideal candidate for drug development and showcase the power of our drug target prediction method in expanding the druggable genome.

**Table 2.**
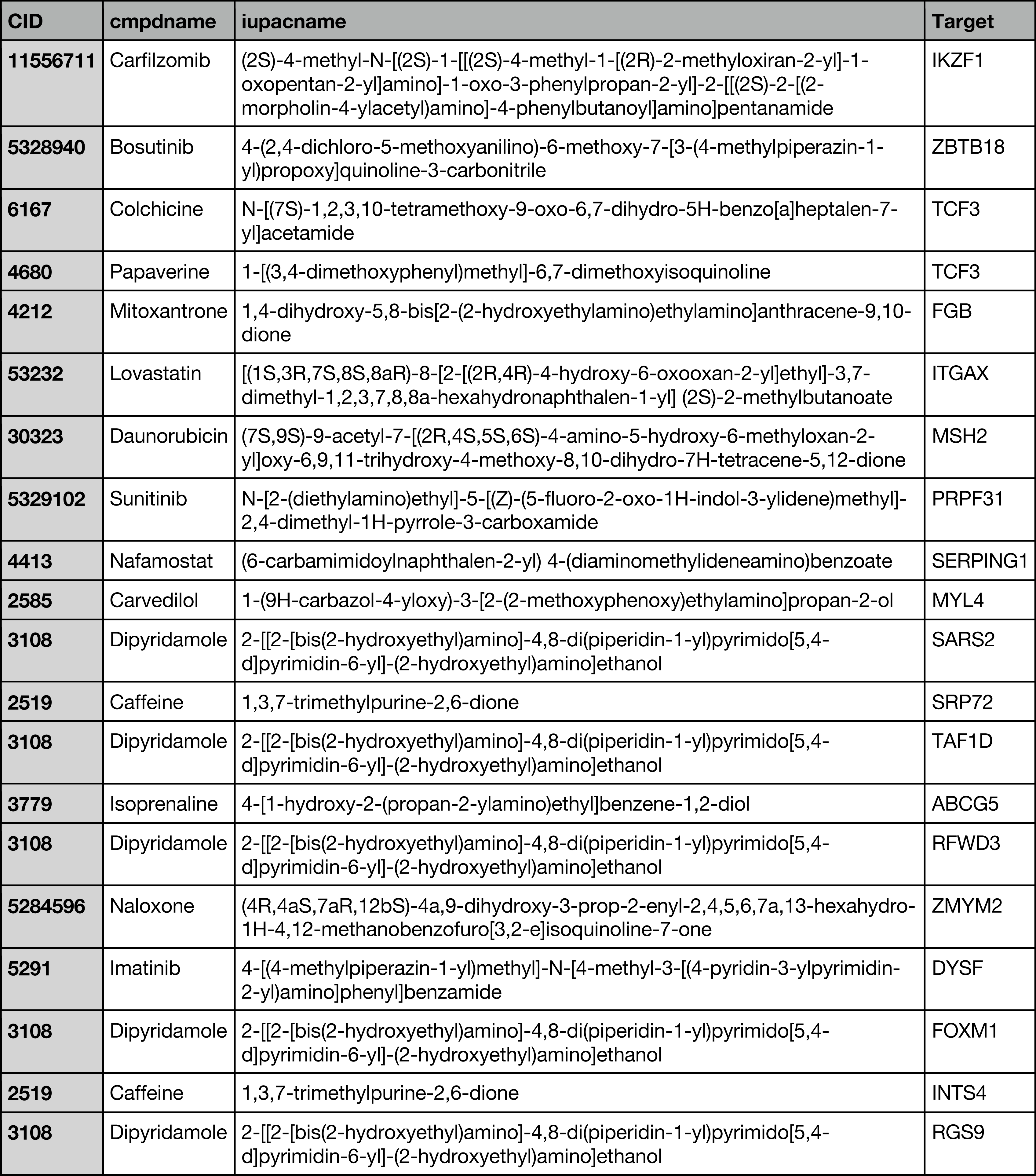
Top 20 drug-target predictions between drugs with known targets and genes that are not yet known to be druggable.

## DISCUSSION

In this paper, we introduce a machine learning method for drug-target prediction that leverages newly proposed features and a novel training scheme. We demonstrate that the new features, including drug-gene phenotype similarity and gene expression profile similarity features, provide complementary information that other features do not capture and enhance the predictive power of our model. In addition, we show that our novel training scheme warrants robust prediction by preventing overfitting and our model achieves accurate prediction while possessing the ability to predict associations for new drugs or currently “undrugged” genes. By doing so, we predict new drug-gene associations and reveal previously unexplored opportunities for drug repurposing and expansion of the druggable genome.

In a conventional train-test split setting, drugs and genes that show up in both the training set and the test set may cause overfitting through data leakage. Although several papers have proposed alternate splitting schemes and experimental settings, including splitting based on drugs or genes [12], few of them adjust their cross-validation strategy, leading to biased performance estimates. Pahikkala et al. proposed a nested cross-validation technique for the same splitting scheme used here [44], yet its complexity renders it computationally expensive considering the fact that for each fold the training and test feature matrices need to be re-calculated (Supplementary Fig. 3a-b). Our newly proposed hold-out validation scheme achieves the same effect of avoiding overfitting with less computation while integrating well with the hyperparameter optimization method. Along with the ensemble approach to solving the class imbalance problem, it can be readily applied to other biological prediction problems, especially those involving biological networks.

As more chemogenomic and phenotypic information about compounds and genes becomes available, we expect that our method will reach even better performance. Since there is no universal method for drug repurposing and druggable gene discovery in place [5, 7, 45, 46], our drug target prediction method can serve as an intermediate step in the drug discovery pipeline, generating reliable candidates which can then be tested by downstream experimental validation. This could greatly accelerate the drug development process and create new opportunities for disease treatment.

## Acknowledgments

The authors would like to thank G. Hooker, J. F. Beltrán, D. Xiong, S. Qian and S. Chen for helpful discussions. This work was supported by National Institute of General Medical Sciences grants (R01 GM124559, R01 GM125639) and National Science Foundation grant (DBI-1661380) to H.Y.

## Author Contributions

H.Y. conceived the study and oversaw all aspects of the study. S.L. performed computational analyses and wrote the manuscript with input from H.Y. S.L. generated figures and tables with input from H.Y. All authors edited and approved of the final manuscript.

## Figure Legends

**Supplementary Figure 1.** (a) Number of known targeters of the gene for all drug-gene pairs. (b) Spearman correlation coefficients of 2D chemical similarity and 3D chemical similarity features. (c) Number of known targets of the drug for all drug-gene pairs. (Statistical significance determined by the two-sided Mann-Whitney U test)

**Supplementary Figure 2.** (a) Schematics of calculating drug-drug similarity features considering known targeters of protein-protein interaction partners of the target in question. (b) Schematics of calculating drug-gene phenotype similarity features considering protein-protein interaction partners of the target in question. (c) Distribution of 2D chemical similarity features considering known targeters of protein-protein interaction partners of the target in question (feature group B). (d) Distribution of side effect similarity features considering known targeters of protein-protein interaction partners of the target in question (feature group D). (e) Distribution of 3D chemical similarity features considering known targeters of protein-protein interaction partners of the target in question (feature group H). (f) Distribution of drug-gene phenotype similarity features considering protein-protein interaction partners of the target in question (feature group F). (Statistical significance determined by the two-sided Mann-Whitney U test)

**Supplementary Figure 3.** Feature matrix calculation. (a) When calculating features for the training set (or fitting the classifier), only associations (drug-gene associations and protein-protein interactions) among train drugs and train targets are visible. (b) When calculating features for the test set (or validation set), all associations are visible except drug-gene associations between test (or validation) drugs and test (or validation) targets.

**Supplementary Figure 4.** Hyperparameter optimization. (a) Change in maximum average AUPR (over 15 train-validation splits) encountered during 1,000 iterations of TPE for each classifier. Dashed lines represent performance using default hyperparameters. (b) Change in maximum average AUROC (over 15 train-validation splits) encountered during 1,000 iterations of TPE for each classifier. Dashed lines represent performance using default hyperparameters.

**Supplementary Table.** All 9,958 novel drug-target predictions above a precision cutoff of 10%.

## METHODS

### Data collection

We collected known drug-gene associations from Probes & Drugs (version 10.2018) [21] along with the SMILES strings of drugs. Side effect were obtained from SIDER 4.1 [47] and OFFSIDES [48], both of which use UMLS concept IDs as identifiers of side effects. However, as similar side effect terms could cause biases in calculating side effect similarity, we mapped all UMLS concept IDs to MedDRA concept IDs using the 2017AB release of UMLS [49]. This allowed us to map UMLS concept IDs to a specific level, PT, of the MedDRA hierarchy, which was obtained from MedDRA (version 21.0) [50]. Disease phenotypes of genes were obtained from DisGeNET (version 5.0) [51]. Similar to side effects, disease phenotypes were also mapped to the PT level of the MedDRA hierarchy, allowing direct comparison with side effects. We only considered drugs with available side effect information and at least one human gene target whose disease phenotypes are known. This resulted in a final set of 11,556 drug-gene associations involving 1,262 drugs and 1,062 human genes. Non-associated drug-gene pairs were obtained by taking all drug-gene combinations not known to be associated using these sets of drugs and genes.

### Feature extraction and feature selection

We extracted 9 groups of features involving 5 feature types: 2D chemical similarity between drugs, side-effect similarity between drugs, 3D chemical similarity between drugs, drug-gene phenotype similarity and expression profile similarity between genes.

For each type of drug-drug similarity feature, we calculated two groups of features: (1) similarity between the drug in question and the known targeters of the gene in question (Fig. 1a); (2) similarity between the drug in question and the known targeters of the protein interactors of the gene in question (Supplementary Fig. 2a). For the former, four different aggregation functions were applied for each group: minimum, mean, median and maximum, while for the latter, mean was replaced by first applying the mean to each set of known targeters corresponding to a single protein interactor before applying a second mean function to obtain a single value. Chemical similarity (both 2D and 3D) was calculated by taking the Jaccard index of the fingerprints of the drugs. 2D (Morgan) fingerprints of compounds were generated with the RDKit Python package, while 3D fingerprints were generated with the E3FP package in Python [23]. Side effect similarity was calculated by taking the Jaccard index of the sets of side effects of the drugs.

Drug-gene similarity was calculated as the Jaccard index between the set of side effects of the drug in question and the set of disease phenotype of the gene in question. We also considered the protein interactors of the gene in question and calculated their closedness to our drug in question in terms of phenotype. Using minimum, mean, median and maximum as aggregation functions we obtained a group of 4 features (Supplementary Fig. 2b). Gene-gene expression similarity was obtained by extracting the expression levels of the two genes from GTEx V7 [24] and calculating their Spearman correlation coefficient across all tissues. Applying the same 4 aggregation functions resulted in another group of four features (Fig. 2c). In total, we extracted a total of 33 features across 9 groups for each drug-gene pair.

Since the features utilize the whole drug-gene association network as well as the protein-protein interaction network, care needs to be taken when calculating features for the training/validation/test sets. When calculating the feature matrix for the training set, only drugs and genes used for training and connections among them were visible (Supplementary Fig. 3a). On the other hand, when calculating the feature matrix for the test set, all associations except those between test drugs and test targets were visible (Supplementary Fig. 3b). This also applied to training and validation sets during hold-out validation, where the full training set can be seen as the full data and the validation set can be regarded as the test set.

To obtain the optimal feature combination, we calculated all features for the training set and applied group maximum concave penalty (MCP) [25] with the ‘grpreg’ R package for feature selection using default parameters. All subsequent training was done with this optimal set of features.

### The training scheme and hyperparameter optimization

We randomly split all drugs into “train drugs” and “test drugs” with a 2:1 ratio, and similarly we split all genes into “train targets” and “test targets” with the same ratio. The training set consisted of all drug-gene pairs where the drug is a “train drug” and the target is a “train target”; the test set consisted of all drug-gene pairs where the drug is a “test drug” and the target is a “test target” (Fig. 3a). To address the class imbalance problem, we split all the drug-gene pairs with negative labels into subsets, each having a size 5 times that of all the positive labels in the training set expect the last one (the size of the last one is between 5 to 10 times that of all the positive labels in the training set). Each subset of negative labels was combined with all the instances with positive labels in the training set to obtain a training subset (Fig. 3b).

For each training subset, we trained an XGBoost classifier [26]. XGBoost is a decision tree ensemble model that additively trains decision trees which predict the prediction error of the existing ensemble. It was chosen for its speed and ability to automatically learn branch directions for missing values. In each classifier the scale_pos_weight hyperparameter was set to the negative-to-positive class ratio in the corresponding training subset. At the end, we applied an ensemble approach by taking the average prediction score of all the classifiers trained as the final prediction score.

To find the best sets of hyperparameters for each classifier, we adopted the tree-structured Parzen estimator (TPE) approach [29]. Instead of cross-validation, we split the entire training set into training and validation sets using the same splitting method as the train-test split to ensure that there was no overlap between data used for training and validation in terms of either drugs or genes (Fig. 3c). For each classifier, the part used for training was then intersected with the corresponding training subset before used as input to the classifier, while the entire validation set was used for performance evaluation. For each classifier and each set of hyperparameter, the split was conducted 15 times, and we selected 1 minus the average AUPR of the 15 trials of hold-out validation as the loss function to minimize for TPE (Fig. 3d). We ran TPE for 1,000 iterations to obtain the best set of hyperparameters that minimized the loss function for each classifier in our ensemble. After obtaining the optimal sets of hyperparameters, we retrained each classifier using all data from the corresponding training subset.

### Model evaluation and application

Model performance on the training set was evaluated by taking the average performance across all classifiers in the ensemble using performance metrics calculated from the best sets of hyperparameters during hyperparameter optimization, including average AUROC and average AUPR across 15 splits. To evaluate the contribution of each type of feature to model performance, we dropped each type of features and repeated training and hyperparameter optimization procedures. To further evaluate the predictive power of our model, we predicted on the left-out test set which had no overlap with training data in terms of either drugs or genes. Predictions were ranked according to their prediction scores to produce the ROC curve and the precision-recall curve. To apply our model to predicting new drug targets, we considered all drug-gene pairs that were not previously known to be associated where the drug had known side effects and the gene had known disease phenotypes. 9,958 new drug-gene associations were identified with a precision lower bound cutoff at 10% (Supplementary Table).

