## Supplementary figures and images for "Revealing new therapeutic opportunities through drug target prediction via class imbalance-tolerant machine learning"

### Supplementary Figures

#### Supplementary Figure 1

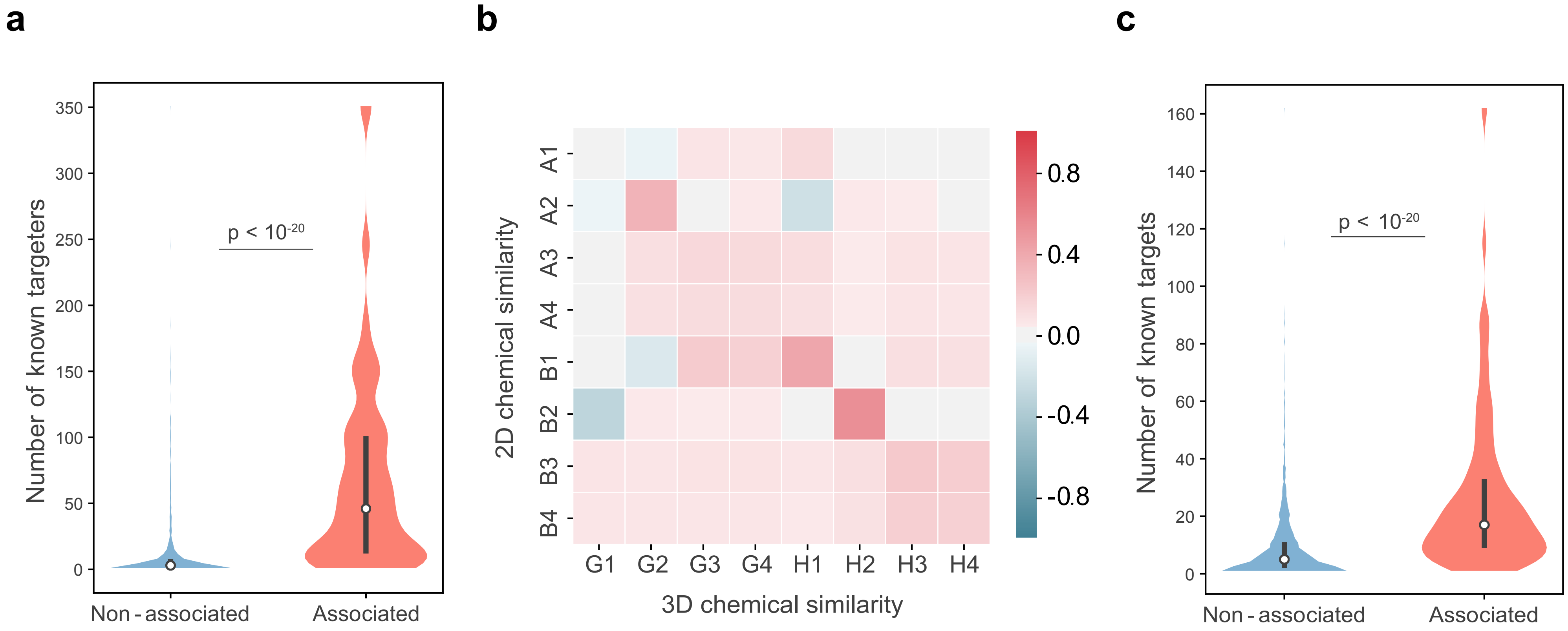

#### Supplementary Figure 2

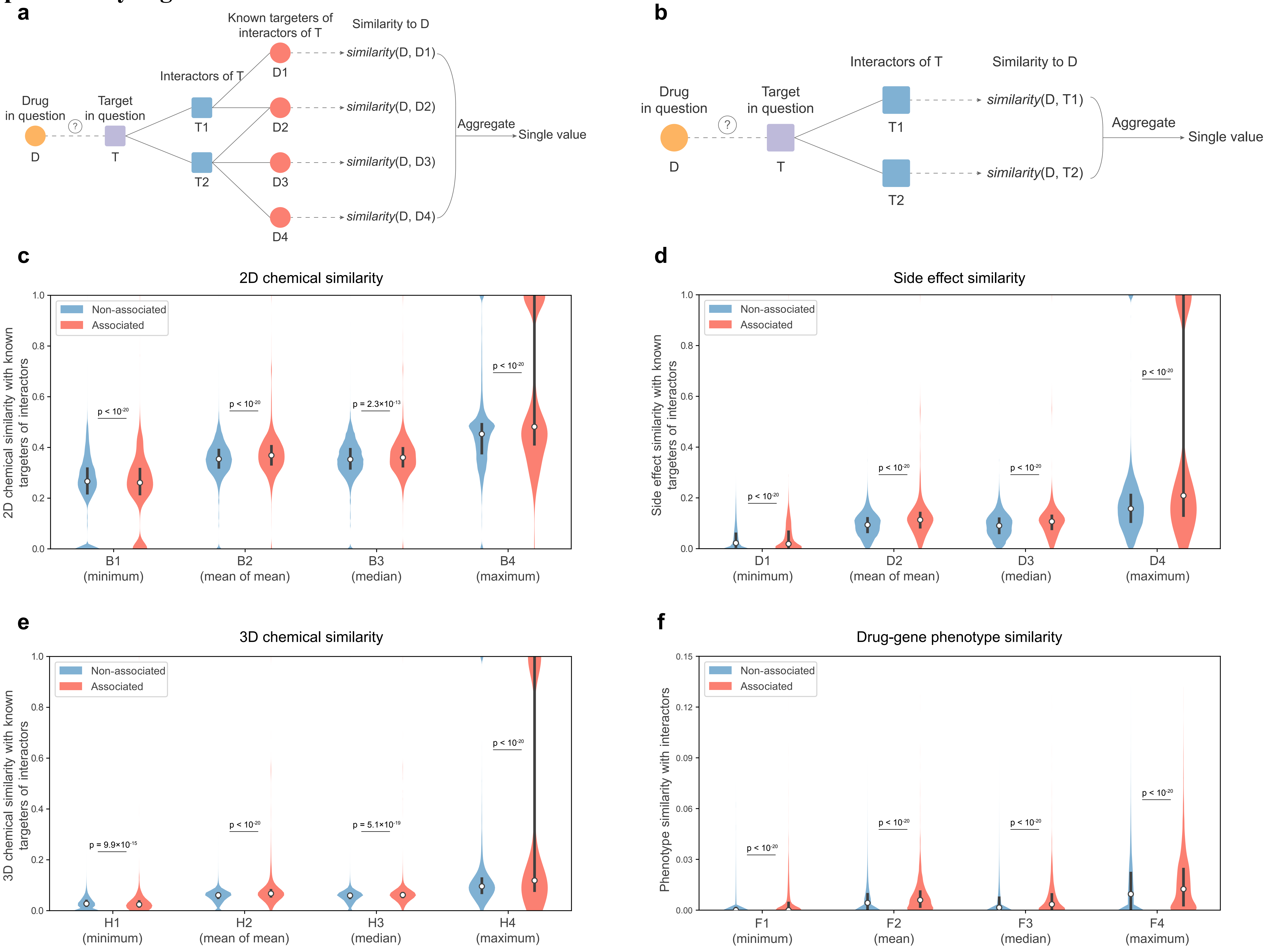

**Supplementary Figure 3**

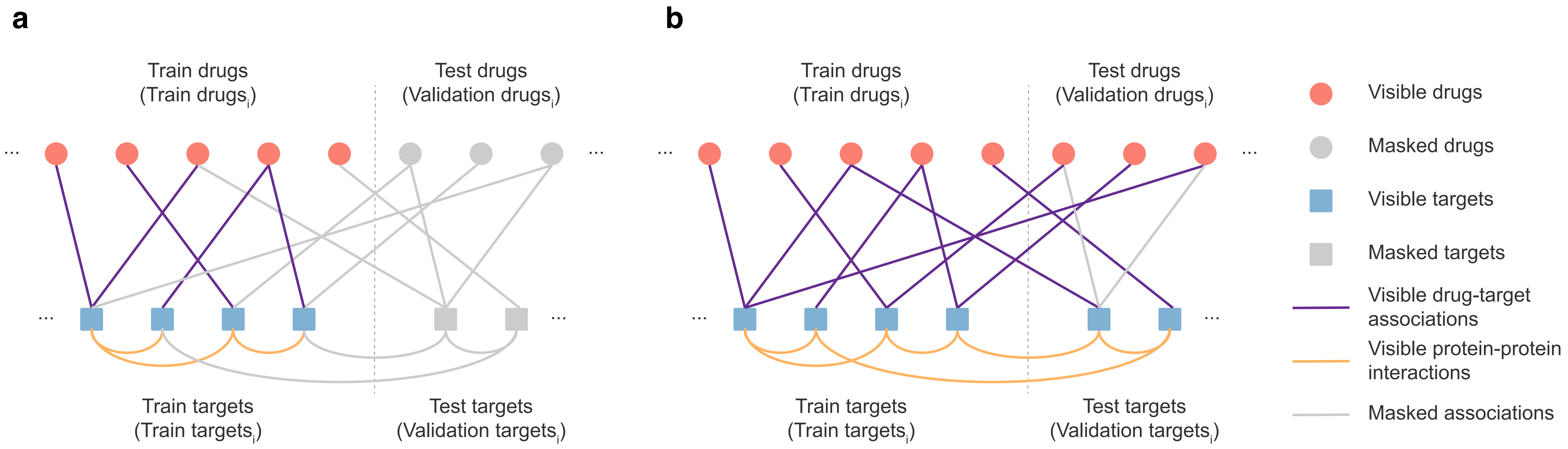

#### Supplementary Figure 4

**a**

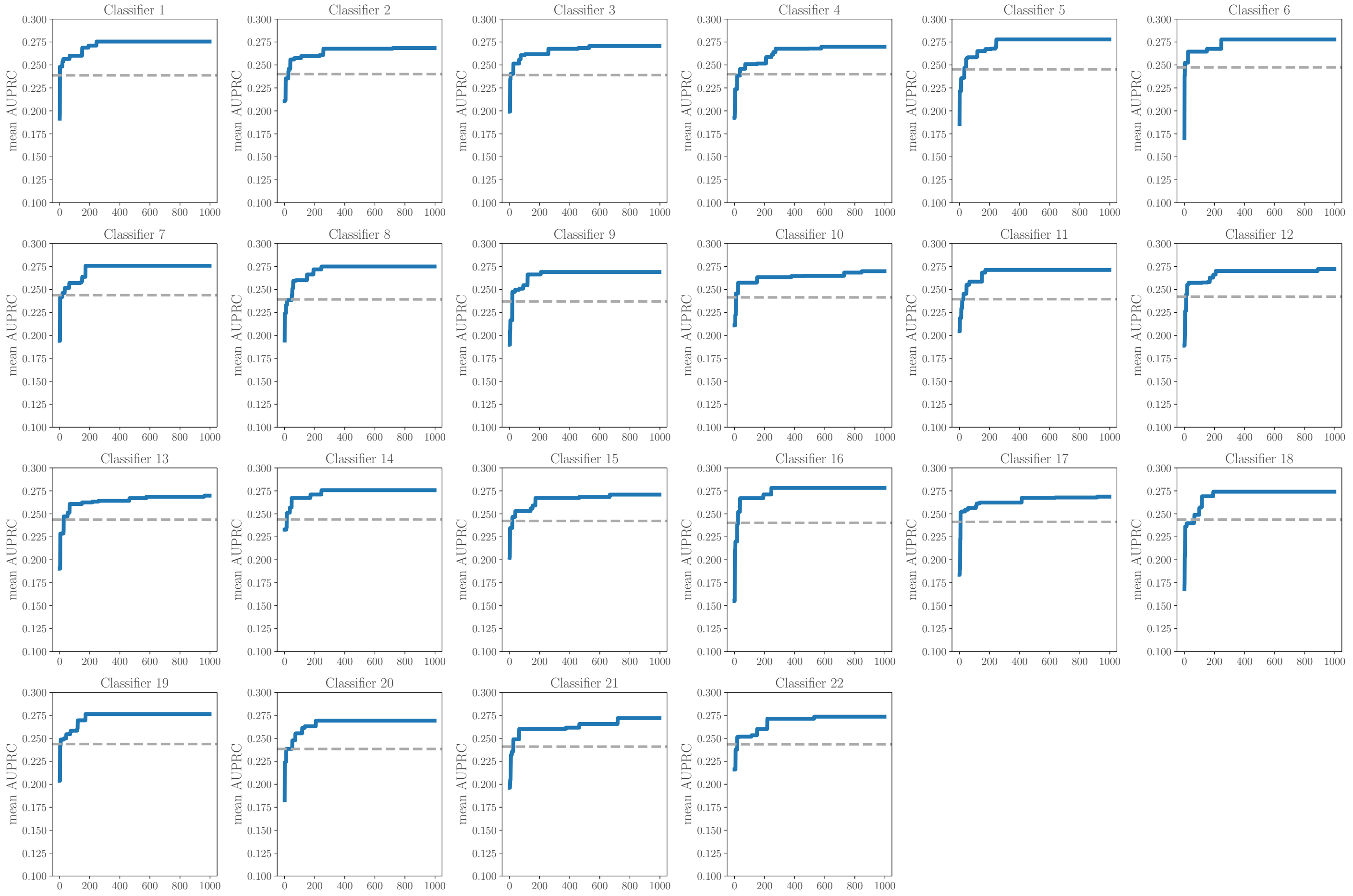


**b**

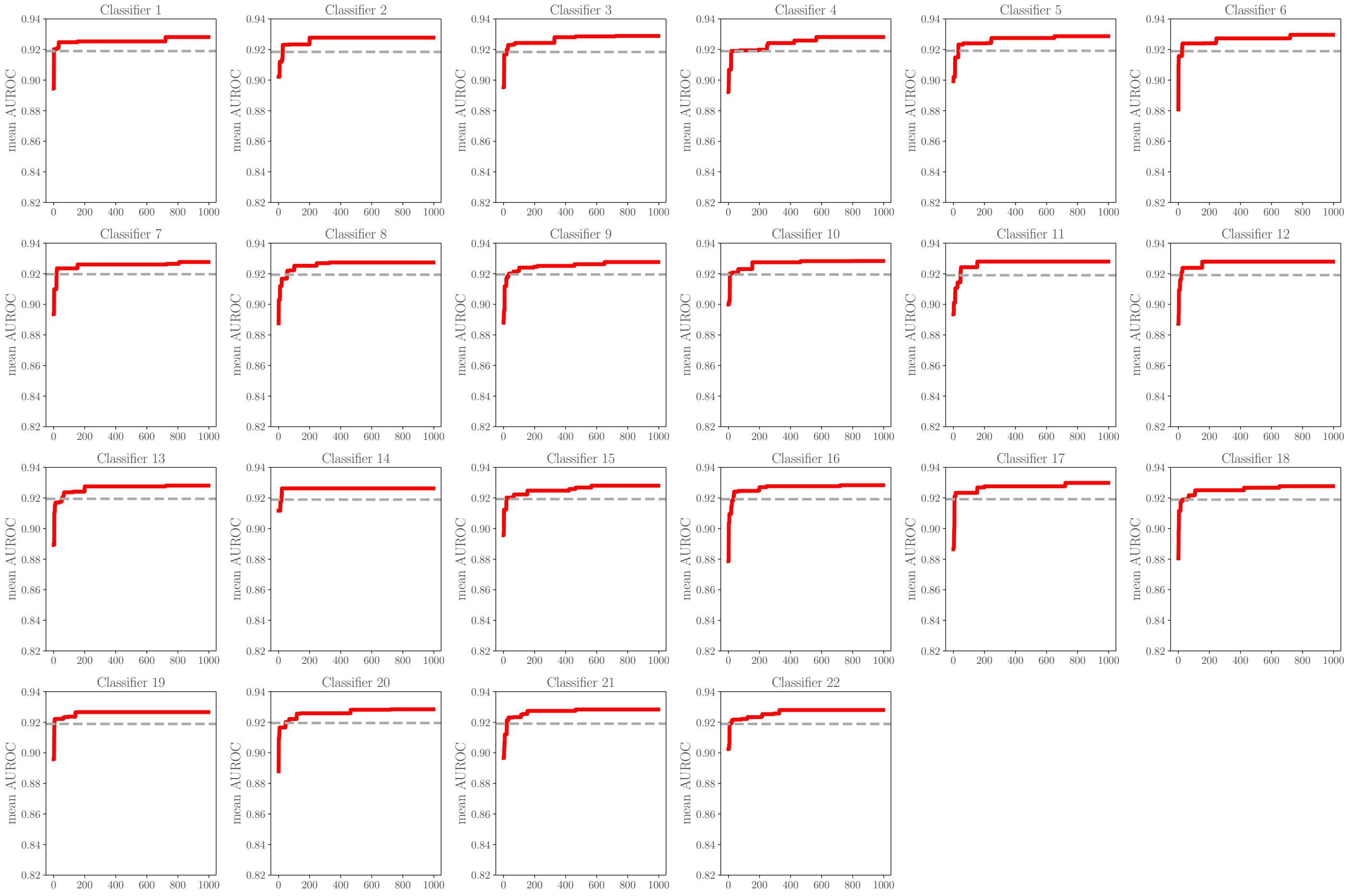
